# Group dynamics and habitat use of the Giant Otter, *Pteronura brasiliensis* (Zimmermann, 1780), in seasonally flooded forest in the Araguaia River, Central Brazil: A 10-years study

**DOI:** 10.1101/2022.01.30.478386

**Authors:** Benaya Leles, George Georgiadis, Thais Susana, Nils Kaczmarek, Reuber Brandão, Silvana Campello

## Abstract

We carried out monthly surveys of the giant otter population between 2010 and 2020 in a study area comprised of 1,500 hectares of igapó flooded forest with oxbow lakes in the Cantão region of central Brazil. We recorded 16-32 resident adults in the study area each year, distributed in 4-8 groups. Resident groups exhibited extensive home range overlap, with each group using several lakes and larger lakes used in rotation by up to six groups. Dens and campsites were also shared by multiple groups, but lakes were used by only one group at a time, and encounters between groups were very rare. 24 adult otters were observed to join an existing group. Some individuals changed groups multiple times. Resident adult turnover was high. Each year an average of 36% of resident adults were new immigrants, and 72% of groups left the area within two years. Resident groups had, on average, one litter every three years, and annual cub production showed high variability and a negative correlation to the number of new immigrants in the area. No pairs of giant otters reproduced successfully during the study. Groups of three otters formed through the recruitment of an adult individual by an existing pair and reproduced as successfully as larger groups. Group dynamics and territorial behavior in the Cantão flooded forest ecosystem, where optimal giant otter habitat is continuous in all directions, were found to be different from that reported in areas composed of patchy (isolated oxbow lakes) or linear (rivers) habitat. This suggest that giant otter social and territorial behavior is plastic and adapts to the spatial characteristics of the habitat.

## Introduction

The giant otter (*Pteronura brasiliensis*) is an endangered top predator of tropical South American lakes and rivers [1–5]. Giant otters originally ranged broadly from the Andes to the coast of Brazil, but they were extirpated from much of their range by hunting for the pelt trade, primarily between 1940 and 1980. Today the easternmost remnant population of the species occurs in the Araguaia river basin of central Brazil.

### The Cantão Ecosystem

The Cantão wetlands ecosystem is located at the confluence of the Javaés and Araguaia rivers, in the state of Tocantins in central Brazil (Fig 1). The region is a sharp ecotone between the Cerrado and Amazon biomes, with exceptional biodiversity [6]. The Javaés, a large black water river, is a 400-km offshoot of the Araguaia that flows around the world’s largest freshwater island, Ilha do Bananal. Where it flows back into the Araguaia it forms a 100,000 hectare inland delta named Cantão, an elongated triangular floodplain crisscrossed by meandering channels and dotted with over 900 oxbow lakes. This is the largest expanse of suitable habitat for giant otters in the Araguaia river basin, and the species is reported to be common in the area [7–8].

**Fig 1.**
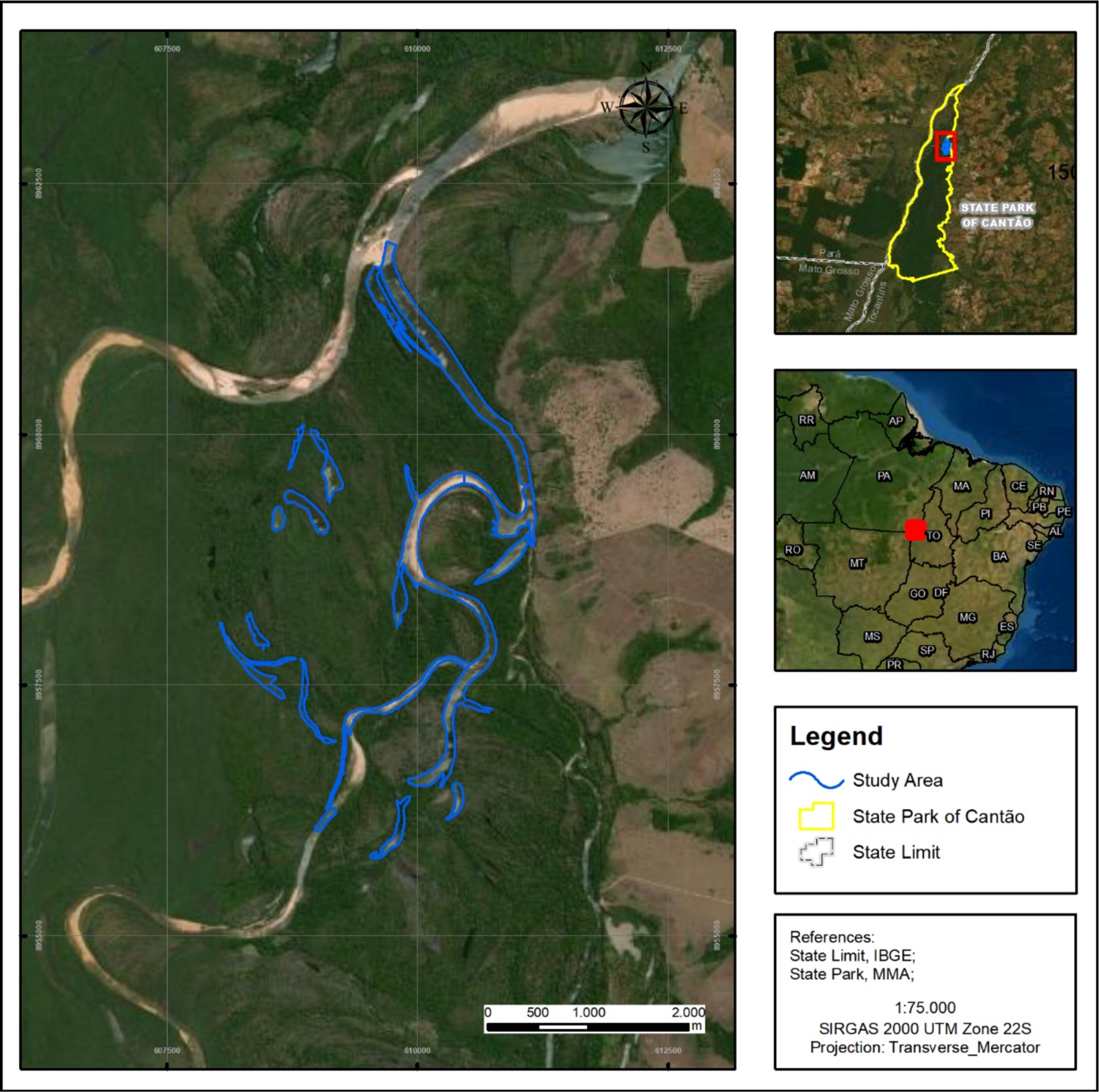
Study area in Cantão, Brazil.

Between December and May, the rising waters of the Araguaia dam the Javaés, and the entire delta floods with dark acidic waters, connecting the lakes. The igapó flooded forest which grows in Cantão is adapted to this cycle, with most tree species growing and fruiting during the peak of the flood and dropping their fruit into the water, where they are consumed by a wide variety of frugivorous fish. The dominant tall tree species are the landi (*Callophylum brasiliensis*) and the piranheira (*Piranhea trifoliolata*), which grow to over 20 meters height. Non-forest habitat includes areas of shrubby vegetation characterized by sarã (*Sapium haematospermum*) and goiabinha (*Psidium riparium*), which turn into marshes where blatterwort (*Utricularia* sp.) and other floating vegetation proliferates in the wet season. Shrub and marsh habitats occur on recently deposited sediment and cover less than 5% of the area of Cantão, but are sunlit and very productive during the floods, concentrating schools of fish like pacús (*Myloplus* sp.) and piranhas (*Serrasalmus* spp.).

In May water levels begin to drop quickly, and between June and September there is little to no precipitation. During the dry season the marshes and flooded forest dry out completely, and fish become concentrated in the lakes and in deep pools along river channels. Most fish predators, including giant otters, arapaima, peacock bass, caimans, and wading birds reproduce during this season.

To the east the Cantão floodplain is bordered by rolling plains of Cerrado vegetation, from which it is separated by the narrow Coco River, actually the easternmost channel of the Javaés delta. To the west it is bordered by the Araguaia river, which is up to three km wide here. Due to the very flat nature of the central Araguaia basin, seasonally flooded habitat similar to that found in Cantão also occurs in narrow strips and on river islands for hundreds of kilometers upstream along the Araguaia and Javaés rivers and their tributaries, although much of this has been altered by dams and irrigation projects in recent years.

Due to the abundance of nutrients made available by the annual flood, the aquatic ecosystem of Cantão is exceptionally rich and productive, hosting over 298 species of fish, whose abundance is among the highest known for Amazonia [9]. At the base of the food chain are many species of pacú, which feed primarily on fruit dropping from the flooded vegetation; piranhas, which are omnivorous, eating fish and falling arthropods as well as vegetable matter; and piaus (*Schizodon vittatus*), whose specialized lips allows them to feed on the rich layer of mucus and microorganisms which covers the submerged vegetation. All of these are in turn preyed upon by an abundance of larger predators, including giant otters [10]. Other large aquatic apex predators that are common in Cantão include black caiman (*Melanosuchus niger*), araguaia river dolphins (*Inia araguaiaensis*), and arapaima (*Arapaima gigas*).

Most of the Cantão ecosystem is protected within 90,000-hectare Cantão State Park. The park is bordered by river channels on all sides and contains over 850 oxbow lakes with surface area greater than one ha, and over 240 km of channels meandering through its interior. Until 2017 the park was considered one of the best managed protected areas in the Brazilian Amazon [11] and was relatively well funded and staffed. The park is completely uninhabited except for a small area near the town of Caseara, which has been used by local people for seasonal agriculture since before the creation of the park. Fishing is prohibited inside the park, and most of it is off limits to unauthorized persons. Despite this, much of the park is vulnerable to invasion by fish poachers, who seek high value species like arapaima and tucunaré (*Cichla* spp.) which have been depleted outside the protected area. These poachers set up clandestine camps and fish nearby lakes until they are depleted. They not only reduce availability of fish prey, but also scare away or shoot giant otters, which they blame for declining fish stocks. Policy changes starting in 2019 have weakened park management, with patrols becoming less frequent and poacher activity intensifying.

## Materials and Methods

We conducted our studies in the vicinity of Instituto Araguaia’s research station. The station is located in Instituto Araguaia’s 540-hectare private inholding within Cantão State Park, thus facilitating logistics for the fieldwork. This area includes 15 oxbow lakes and 9,300 m of river channels. During the low water season, the river channels themselves become a string of long deep pools, ecologically very similar to oxbow lakes, separated by shallow sandbanks. The study site includes one of these stretches of river channel, totaling 16 lakes or lake-like bodies of water which retain water depths greater than two meters during the dry season. The largest lake, Lago Grande, is 2,220 m long and 110 m wide and remains connected to the river channel year-round. The other lakes range from 230 m to 1,218 m in length. These bodies of water are contained within a perimeter encompassing approximately 1,500 hectares of igapó flooded forest, with some marshes. The site is representative of the Cantão ecosystem as a whole, containing roughly proportionate samples of each of the park’s natural communities. It is also one of the sectors of Cantão State Park least impacted by fish poachers, who are dissuaded by the year-round presence of Instituto Araguaia’s researchers, rangers, and volunteers.

Surveys followed the ‘Population Census Methodology Guidelines for the Giant Otter [12]. In the dry season, lakes and river channels were surveyed using canoes, and isolated lakes were surveyed on foot. In the wet season, lakes, channels, flooded marshes, and igapó forests were surveyed by canoe. Traditional dugout canoes, as well as fiberglass canoes, were used, powered by 44-pound electric motors. We found that giant otters are less disturbed when approached with an electric motor than by paddling because with the electric motor the observer can remain silent and motionless.

Surveys were conducted between sunrise and 11:00h, and between 15:00h and sunset, which are the giant otters’ peak hours of activity [2, 13]. When giant otters were sighted, the group was followed from a distance large enough to avoid alarming the animals (between 30 and 200 meters, depending on the behavior of the group). Panasonic DMC-FZ series cameras with 18-50x optical zooms were used to film and photograph the giant otters, allowing subsequent identification of individuals by their throat markings and accurate counts of group size. Data from direct observations were complemented with images from Reconyx and Bushnell camera traps (several models over the years), placed at the entrance of active dens, on campsites, and along giant otter trails between lakes.

Survey years were defined to extend from May 1 to April 30 of the following year to coincide with the period between peak floods and to encompass a full giant otter reproductive season. For survey purposes, two stretches of the river channel within the core area that remain deep enough in the dry season for giant otters to swim and forage were classified as “lakes”. Both of them are isolated from other water bodies by extensive shallow areas during the dry season and are very similar to lakes in terms of dimensions and habitat characteristics.

Regular surveys started in September 2010 encompassing four lakes around Instituto Araguaia’s research station. In 2011 surveys were carried out in eight lakes, in 2012 in 12 lakes, and from 2013 to 2019 surveys covered the entire core study area, defined to include 16 lakes and lake-like stretches of deep river channels. In 2020 it was not possible to adequately survey one group of three lakes on the eastern edge of the study area because armed fish poachers set up a permanent camp in the area during the dry season. Surveys were conducted monthly between August 2010 and April 2021, for periods varying between four and 23 field days. In the dry season (June-November) every lake in the core area was surveyed by researchers at least once a week, with camera traps left at active dens and campsites. In the wet season access to parts of the area was blocked during periods when water levels were too high for surveys on foot but not high enough to allow access by canoe. During these periods most lakes were surveyed at least once a month, and temporarily inaccessible lakes were surveyed using camera traps left at known wet season den sites for periods of up to two months. Additional data was obtained during annual expeditions to lakes in the region adjacent to the core study area, as well as to other sectors of Cantão Park.

A sighting catalog for individual giant otters was developed according to Groenendijk et al. [12]. Individuals were identified by their unique throat markings. Each individual entered into the catalog was given a name and an identifying number. Each group recorded also received a group number. Groups whose composition changed were considered to be the same group when at least 60% of the individual members remained constant [12]. Sex information was obtained when possible by the identification of sexual characteristics in videos and camera trap images.

Giant otters were considered to be resident in the core study area if they exhibited territorial behaviors (denning, use of latrines, or actively approaching intruders while periscoping) and were recorded within the study area on at least three separate days for a period of 30 or more days. Groups and solitary otters that did not fulfill these criteria were considered to be transient and were excluded from the analyses of habitat use and range overlap but were included in the analyses of group size and composition. Giant otter records obtained inside the flooded forest during the wet season were assigned to the nearest lake for home range evaluation.

Animals recorded were classified into three age groups: “Newborn cubs” were defined as animals up to around 60 days old, which remain inside the den and cannot enter the water on their own, although they may sometimes be seen briefly outside the den entrance, or while being carried by adults; “free-swimming cubs” are animals 60-180 days old which are able to enter the water on their own, initially for brief swimming lessons with the adults and later to follow the adults in their daily foraging, and which can be identified as cubs by their swimming behavior and size [12]; all other animals were classified as “adult-sized”.

It was often possible to distinguish juveniles up to one year old from older individuals but to be able to use data from multiple observers, we did not use this information in our analysis. Only records of free-swimming cubs were included in the data analysis. Records of newborn cubs could only be obtained opportunistically and were excluded from the analysis because we had no way of determining the mortality rate of cubs before they became free-swimming and could be observed reliably. Litter sizes and cub survival rates were calculated based on the number of free-swimming cubs recorded each season that survived until the following year.

Annual turnover rate of resident adult individuals at the study site was calculated by dividing the number of resident adults that were not resident on the previous year by the total number of resident adults on the site each year. The same calculation was performed for resident groups. Dispersal distances were calculated both in a straight line and along the shortest water route, following meandering river channels and lakes.

Although we analyzed our results in the light of published material regarding giant otters, we did not identify many long-term continuous surveys similar to our study which could serve as a comparative parameter to our data.

## Results

We obtained 3141 records of giant otters during the study, being 2651 through camera traps and 490 through direct observation. We were able to identify 168 individual giant otters. The total number of adult-sized otters recorded in the studied area each year varied from 16 to 32 (mean = 23; SD = 6), distributed between 4 and 8 groups (mean = 5; SD = 1.2) (Fig 2).

**Fig 2.**
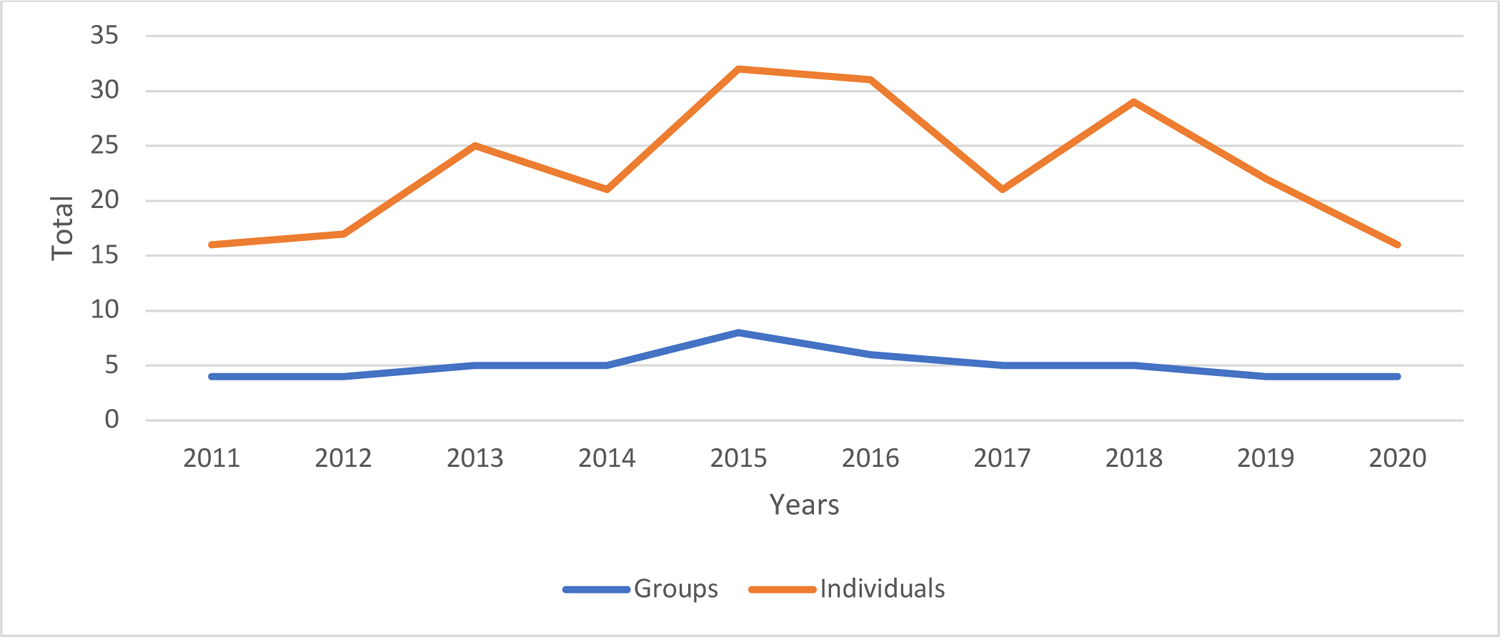
Total number of giant otter individuals and groups in the study area in Cantão, Brazil.

The annual turnover of individuals and groups in the core study area was high. Between one and 17 of the resident adult giant otters recorded each year (or 5-68% of all resident adults recorded; mean = 36%) were new residents that had not been present in the previous year (Fig 3). These immigrants moved into the study area either as entire groups or as individuals that joined a resident group (Fig 4). 72% of groups whose arrival date into the study area was known remained resident in the area for two years or less before moving elsewhere. A single group remained in the area for 8 years and is still present as of July 2021 (Fig 5).

**Fig 3.**
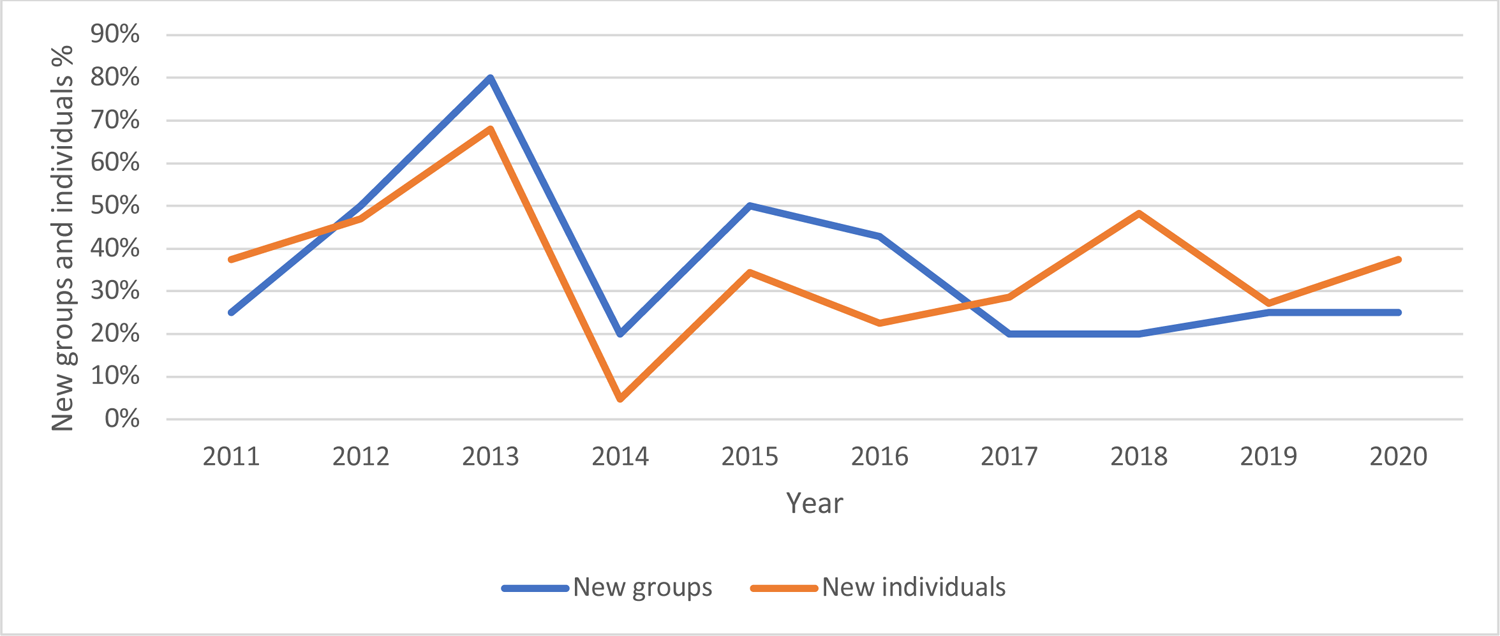
Annual turnover rate of individuals and groups (%) of giant otter in the study area in Cantão, Brazil.

**Fig 4.**
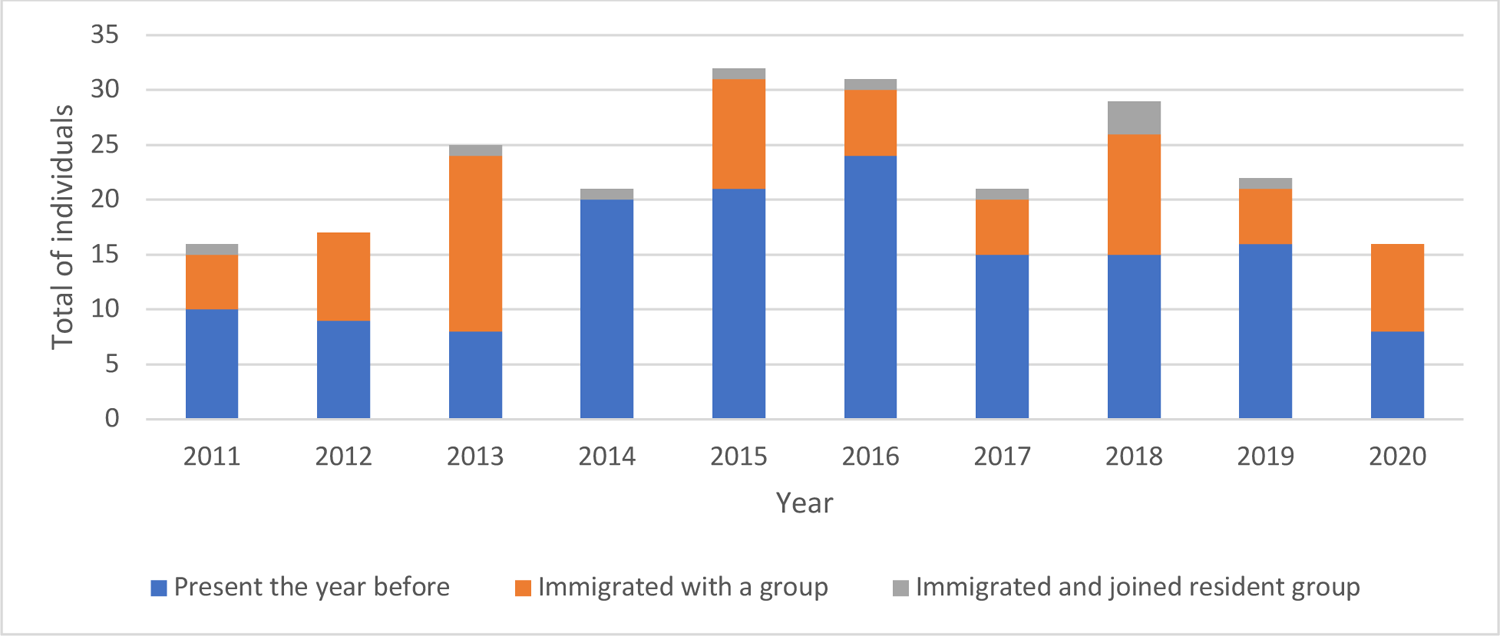
Origin of individuals of giant otter recorded in the study area in Cantão, Brazil.

**Fig 5.**
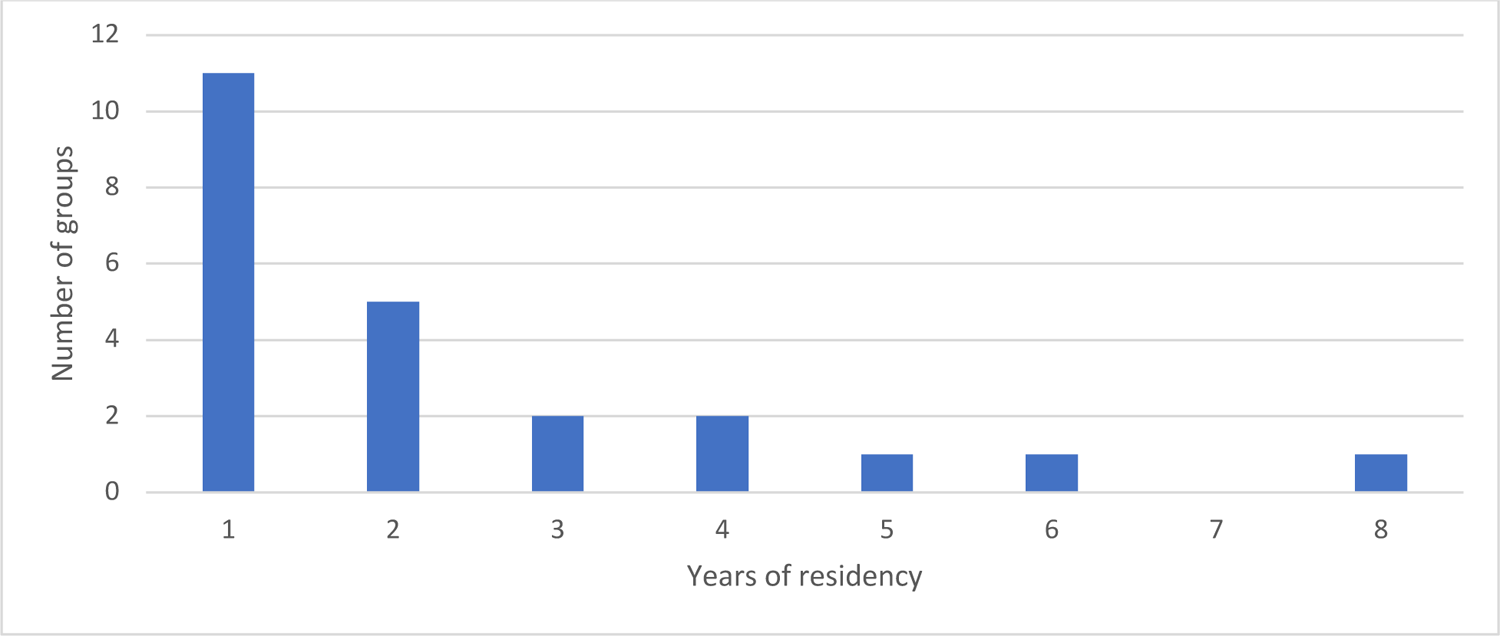
Years of residency of giant otter groups in the study area in Cantão, Brazil.

### Habitat Use and Range Overlap

Resident giant otter groups exhibited extensive home range overlap within the core study area. Most groups used several lakes throughout the year, but the set of lakes used by each group changed from year to year. Small, isolated lakes tended to be used by a single group each year. Over the ten years of the study, in the 11 monitored lakes that are less than 1,000 meters long, only five instances were recorded of a lake being used by more than one group during the same year. Three of these instances took place in lakes located within 100 meters of a larger lake which was also being used by one or more of the groups. Of the 5 monitored lakes longer than 1000 meters, two (Quebra-Linha and Lago do Estirão) are undergoing siltation due to natural processes and are significantly narrower and shallower than most large Cantão lakes. These were used by 1–3 groups each year. The remaining three bodies of water longer than 1000 meters (Lago Grande, Estirão, and Lago das Ariranhas) are relatively wide and deep over most of their lengths (Fig 6). Throughout the survey, most (52 %) records of resident groups were obtained in these three lakes, with each lake being used by up to six groups at different times over a single year.

**Fig 6.**
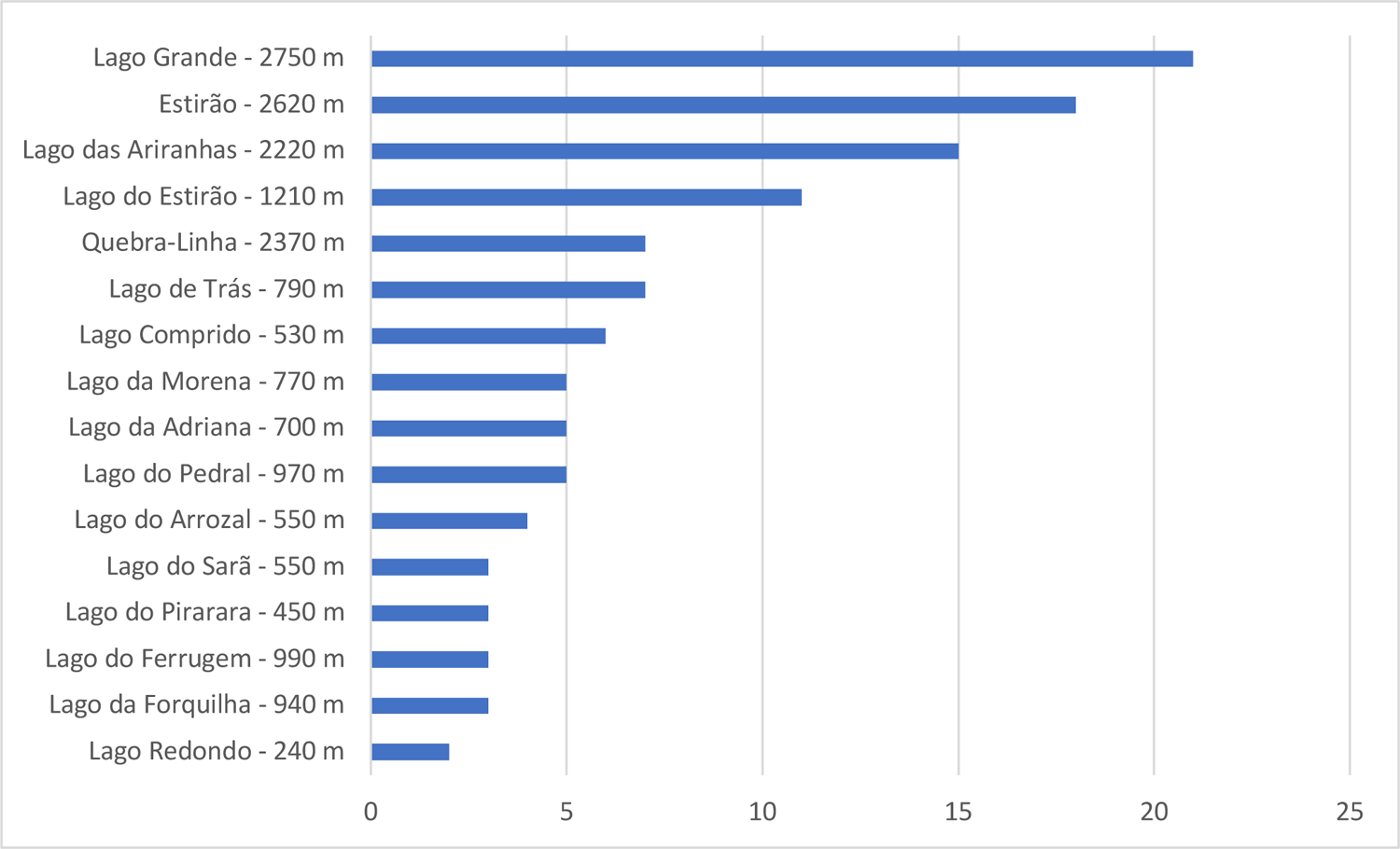
Total number of groups of giant otter recorded at each lake in the study area in Cantão, Brazil.

Lake usage patterns observed were highly variable (Fig 7). Groups rarely remained in each lake for more than a few hours before moving on. A group might use a particular lake once or several times over a few days, and then not return to it for a period ranging from a few days to until the following year. Some groups returned to certain lakes regularly over multiple years, while other groups used them sporadically, with long intervals between visits.

**Fig 7.**
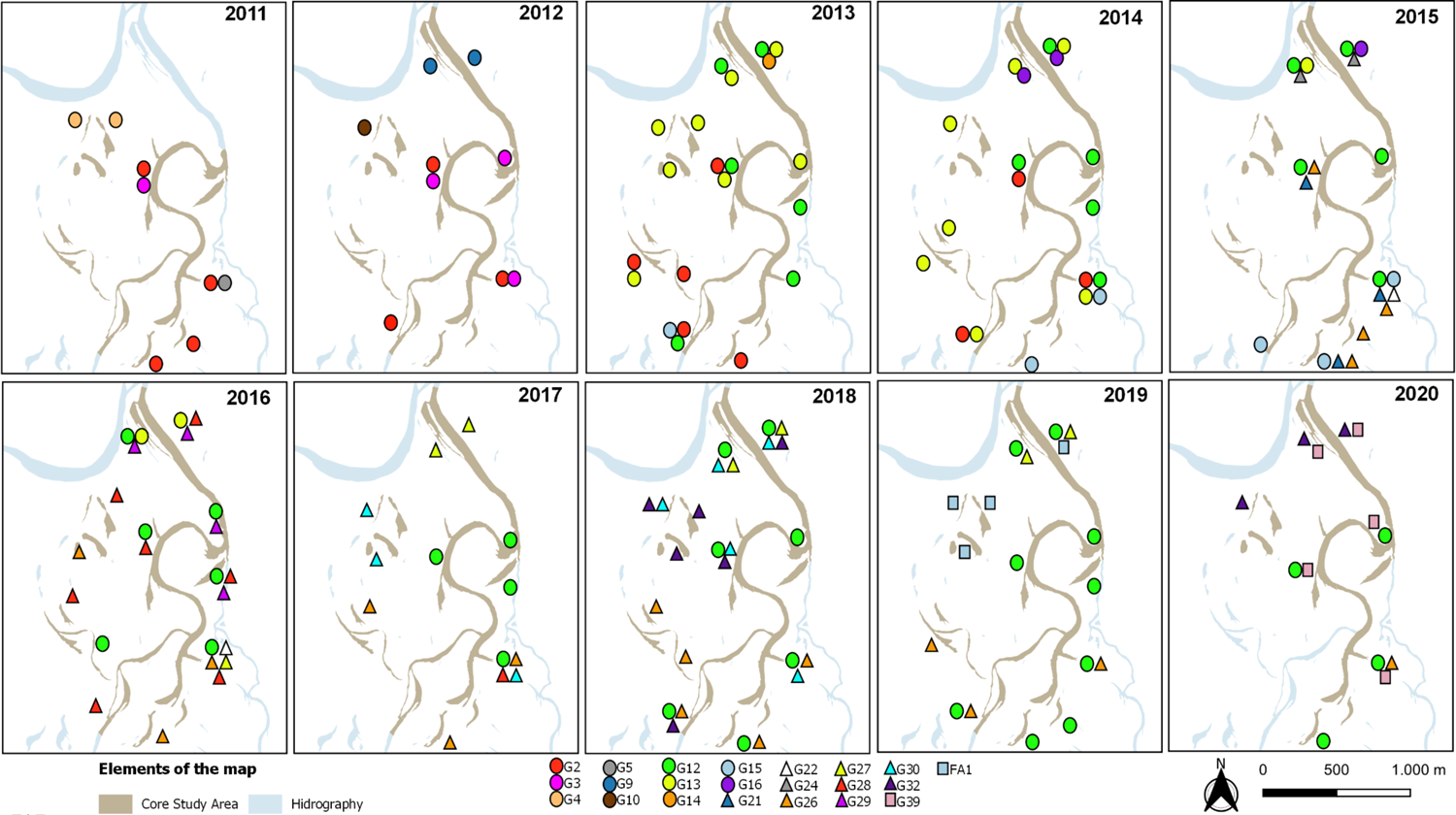
Lake usage by resident groups of giant otter in the study area in Cantão, Brazil.

During periods when two or more groups shared a lake, usage patterns varied from each group using the lake on different months to two or more groups using the lake on alternate days over 1 – 2 weeks. Only 16 times throughout the survey there was more than one group recorded in a given lake on the same day. While using lakes, groups tended to use the same dens and campsites as the previous groups. Denning sites and campsites in prime locations were used almost continually by as many as 11 different groups throughout the survey.

During the annual floods, giant otters extended their range into the flooded areas between lakes. Most giant otter encounters during this season occurred inside the flooded vegetation, although most records of throat markings were obtained while the animals were crossing open water or by camera traps. During the floods, groups were observed to use lakeside dry-season dens and campsites located on ground high enough to remain above water, but also campsites on patches of high ground within the flooded forest, far from open water.

### Group Dynamics

Observed group size ranged from 2 to 8 individuals (mean = 4.1, median = 4, n = 55) (Fig 8). Group size and composition changed over time with births, adult individuals joining existing groups, and individuals disappearing.

**Fig 8.**
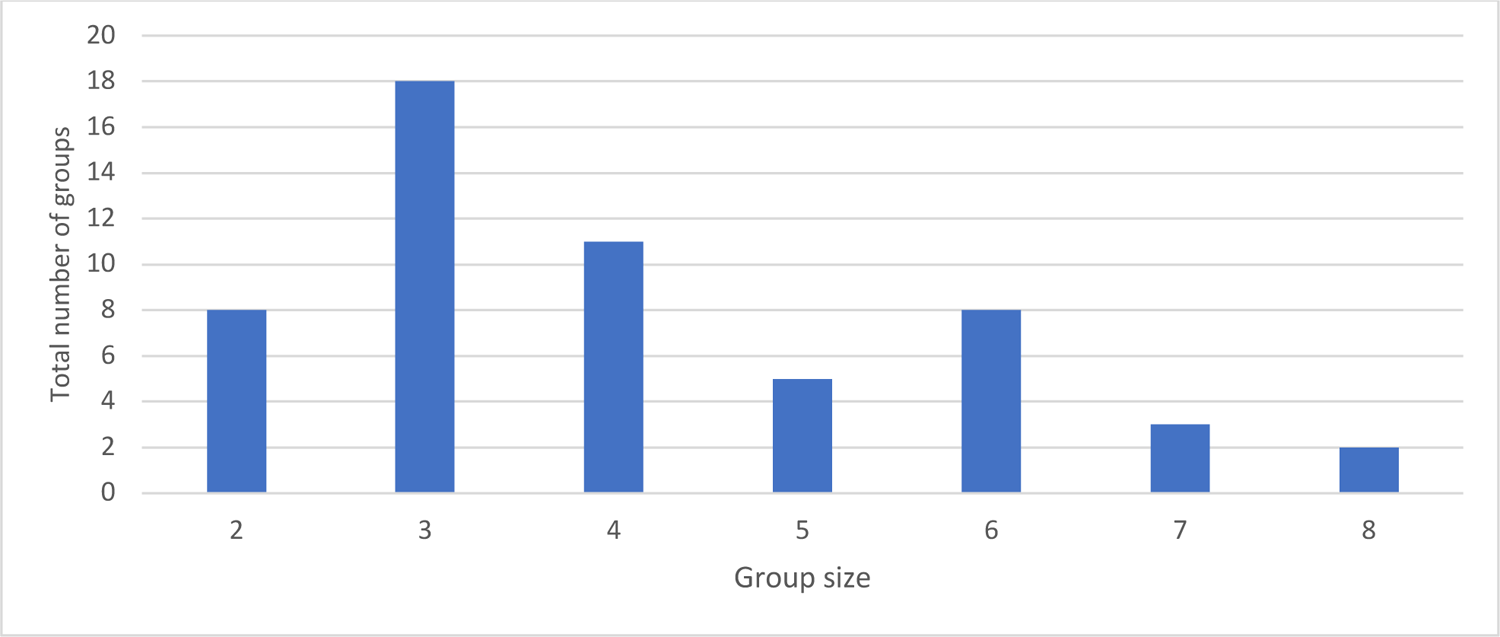
Size of groups of giant otter in the study area in Cantão, Brazil.

We recorded 24 episodes of adult-sized giant otters joining a group of two or more animals. In 11 of these observations, the new otter was formerly a member of a resident group. Three otters joining groups were resident solitaries, and 10 were new to the area. In two cases the new group members subsequently left the group and joined a different group. Of 18 adult-sized individuals of known sex that joined existing groups, 13 (72%) were male. Of the individuals that were new to the study area and joined a resident group whose sex was determined, three were male and two were female. In 16 cases where we were able to determine the status within the group of the new member, three became the reproductive female, three became the reproductive male, and the remainder (two females, two males, and six of undetermined sex) became non-reproductive subordinate group members. One of the subordinate females became the breeding female of her group when the original breeding female disappeared after three years. 19 groups formed by three giant otters were recorded in the core study area. 18 of these trios (94%) were resident groups. Six groups of three otters were formed when a preexisting pair of giant otters was joined by a third adult-sized individual, and one group of three otters was composed of former members of three different resident groups. This group was first recorded after it had formed so it was not possible to determine whether it also started as a pair that was later joined by a third individual. Of the five individuals that joined a pair to form a group of three whose sex was determined, four were male. One group that was first recorded as a trio consisted of two males and one female. The remaining groups of three were either already formed when first recorded or were the remnants of a larger group that had lost members. We observed immigrant groups or individuals of giant otters dispersing distances up to 16.5 km of linear distance (Table 1).

**Table 1.** Dispersal distances for immigrant groups of giant otter and individuals of known origin, in the study area in Cantão, Brazil.

| Individual or Group | Dispersal year | Place of Origin | Dispersal place | Linear distance | Shortest water route |
| --- | --- | --- | --- | --- | --- |
| Pb_52 | 2014 | Estirão | Lago das Ariranhas | 10.92 km | 13 km |
| Pb_52 | 2015 | Lago das Ariranhas | Lago Grande | 14.05 km | 17,4 km |
| Pb_81 | 2015 | Lago da Lua | Lago Grande | 16.5 km | 31,6 km |
| Pb_52 | 2016 | Lago Grande | Paredão | 5.93 km | 9,44 km |
| FA1 | 2019 | Lago do Pequizeiro | Lago Comprido | 9.01 km | 13,06 km |

28 solitary giant otters were recorded in the core study area. Eleven of these met the criteria to be classified as a resident, and three of these were formerly members of resident groups. The remaining solitaries were transients. Resident solitaires were often observed to approach boats and periscope. Nine of the solitary otters recorded subsequently formed pairs or joined existing groups in the study area, and seven of these had been resident solitaries during the previous year. Of 12 solitary giant otters whose sex was determined, eight were male and four were female. Of 4 transient solitaries of known sex, three were male.

23 pairs of giant otters were recorded in the study area throughout the survey. Of these, nine were residents and 14 were transient. Only one resident pair remained in the area as a pair for more than one year. All other resident pairs either left the area or were joined by a third adult animal within one year. 21 pairs were formed during the study. All consisted of at least one member that was a former resident of the study area, and in five pairs both members were former residents. Only two pairs were formed by new members to the area. Of 28 individual otters of known sex that formed pairs, 17 (68%) were former residents (eight males and nine females) and eight were new to the area (six males and two females).

Fig 9 illustrates the changing composition and exchange of members over time of 17 giant otter groups monitored during the study. Group 2, a breeding resident group, was joined by an adult-sized female in 2011, which remained subordinate to the breeding pair. In 2013 the breeding male disappeared and was replaced by Pb 53 a new male arrival. In 2014 this new male bred with the 2011 female and was assisted in rearing the cubs by two remaining offspring of the original breeding pair. In 2015 the group left the core study area. Group 12 immigrated into the study area with three members in 2012, being two males and a female, and was soon joined by a female (Pb 30) whose parental group disappeared from the study area. Within a week one of the males left G12 to become the breeding male of G13, a group of five otters. In 2014 G12 had a litter, and Pb 30 acted as a babysitter for the breeding pair. In 2013 Pb 30 left the group to form a pair with a Pb 91, a new arrival to the study area. G12, now with three adult-sized members, had two more litters, and in 2019 was joined by a male giant otter that assumed a subordinate role to the breeding pair. Meanwhile, after an unsuccessful attempt to breed, individual Pb 30 disappeared from the area and her mate, Pb 91, formed a trio with two other animals, one of which was Pb 53, which had left G2 and returned to the study area after a year’s absence.

**Fig 9.**
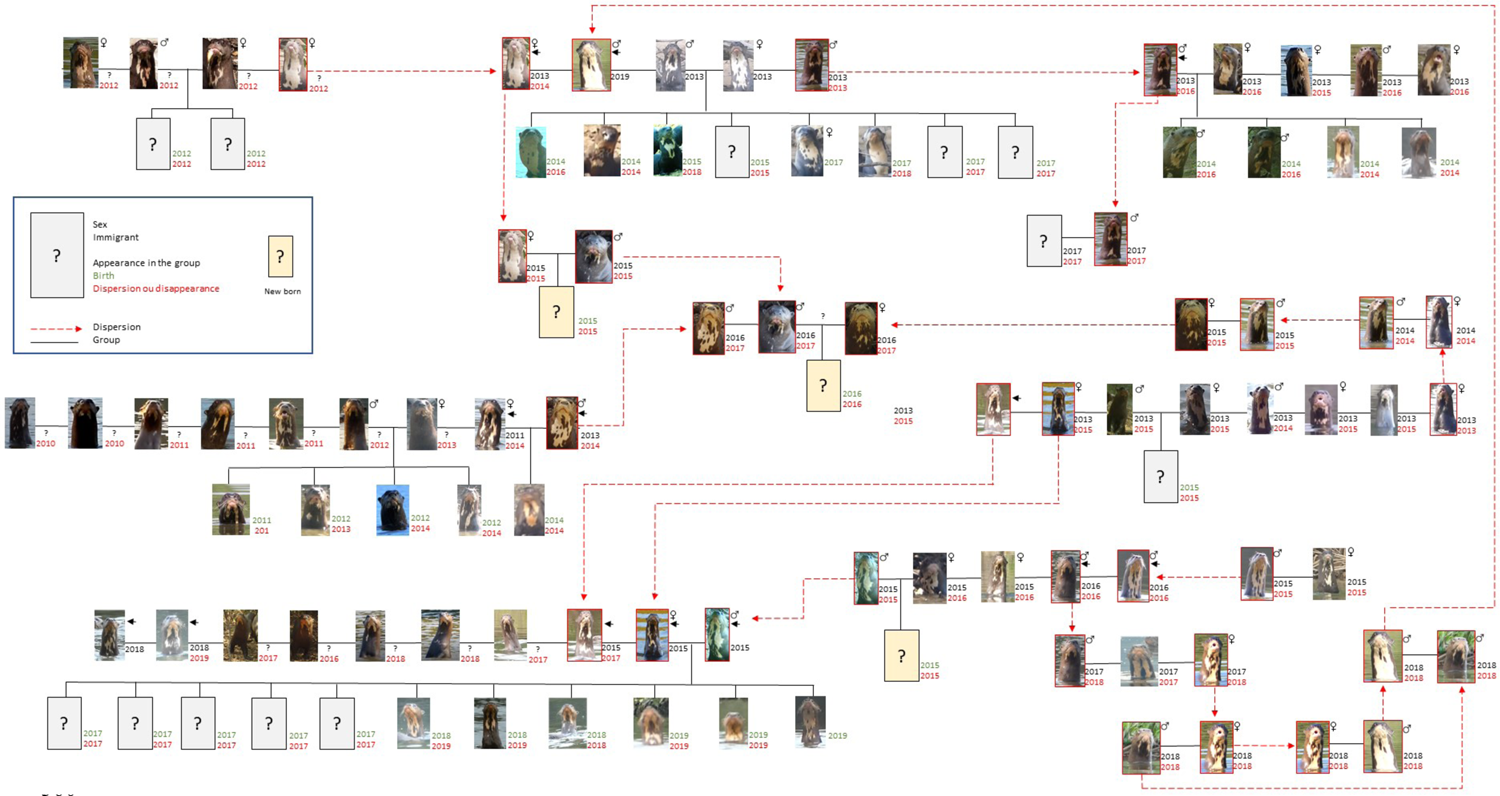
Member turnover and formation dynamics of giant otter groups in the study area in Cantão, Brazil.

### Reproduction

We recorded 17 litter events that produced 42 cubs that reached the free-swimming stage. Litter size at the free-swimming stage ranged from 1-5 (mean = 2.5; median = 2) (Fig 10). Seven cubs too young to enter the water on their own were also recorded, and five of these (71,5%) disappeared before reaching the free-swimming stage. All first records of free-swimming cubs occurred between June and December, suggesting that births took place between April and October.

**Fig 10.**
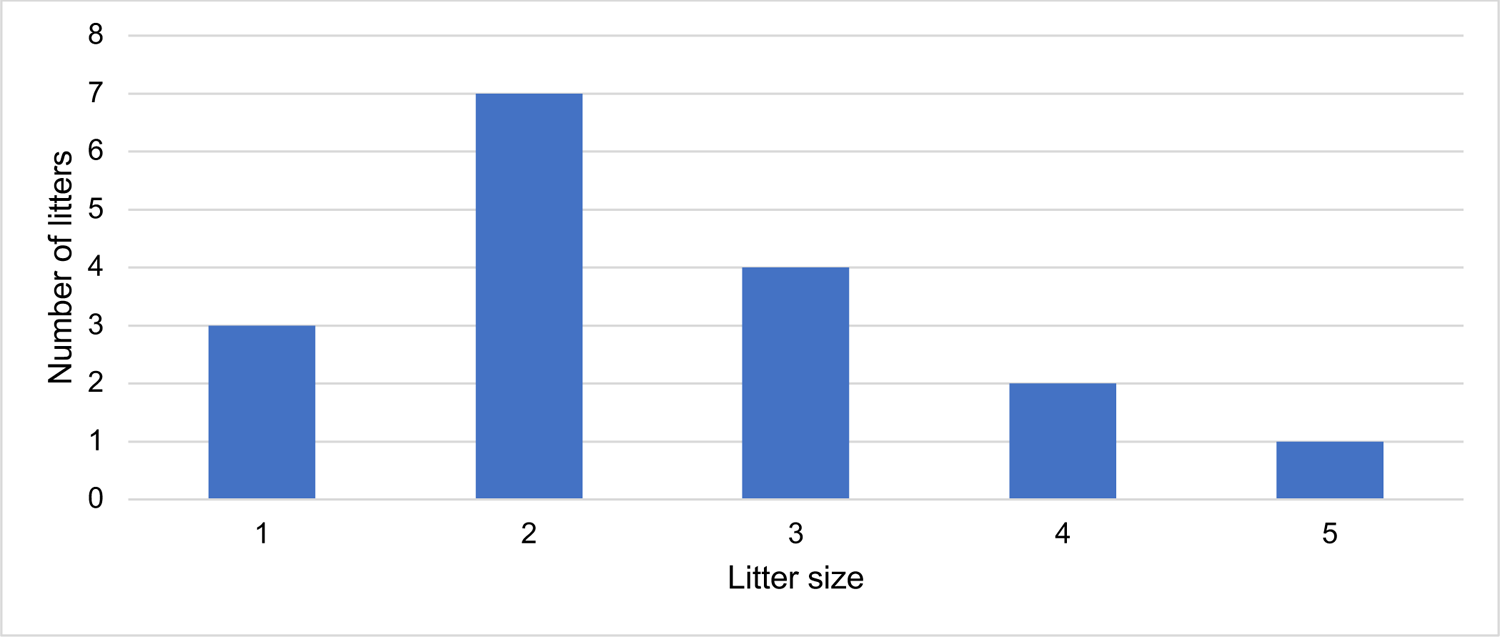
Number of litters and litter size of giant otters in the study area in Cantão, Brazil.

The number of cubs produced per year in the core study area between 2012 and 2020 ranged from 0 to 12 (mean = 4.2) and showed high annual variability (Fig 11). Resident groups had, on average, one litter for every three years of residency (N = 55). The average number of free-swimming cubs produced per year per resident adult (including both breeding and non-breeding group members) was 0.18 (N = 228).

**Fig 11.**
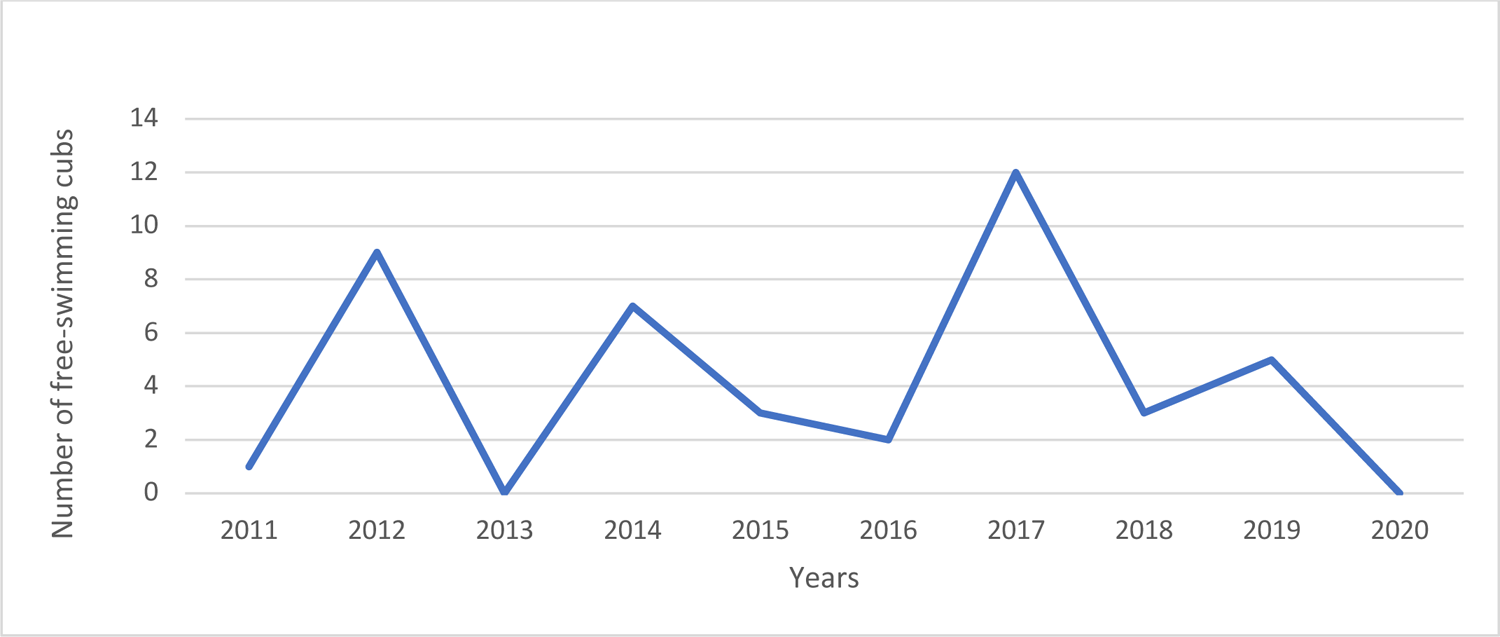
Annual number of free-swimming cubs (approx. 60 days old) of giant otters in the study area in Cantão, Brazil.

Only one pair was observed to have reared cubs to the free-swimming stage while residing in the core study area, and these cubs disappeared a few weeks later. Two other pairs were recorded to have produced newborn cubs that did not survive to the free-swimming stage (Table 2).

**Table 2.** Number of litters by group size of giant otter in the study area in Cantão, Brazil.

| Group size | Number of group-years | Number of litters | Litters per group-year | Total free-swimming cubs | Cubs/group | Free-swimming cubs/adult (all adults) |
| --- | --- | --- | --- | --- | --- | --- |
| 2 | 8 | 1 | 0,13 | 1 | 0,13 | 0,06 |
| 3 | 18 | 7 | 0,39 | 16 | 0,89 | 0,30 |
| 4 | 11 | 3 | 0,27 | 5 | 0,45 | 0,11 |
| 5 | 5 | 1 | 0,20 | 4 | 0,80 | 0,16 |
| 6 | 8 | 5 | 0,63 | 16 | 2,00 | 0,33 |
| 7 | 3 | 0 | 0,00 | 0 | 0,00 | 0,00 |
| 8 | 2 | 0 | 0,00 | 0 | 0,00 | 0,00 |
| **Total adults**<br>**= 210** | 55 | 17 | 0,31 | 42 |  | 0,2 |

The mean number of free-swimming cubs born per resident group year (including years when a resident group had no free-swimming cubs) was 0.76 (n = 55 group years). Pairs of giant otters had the lowest number of litters per group-year, while groups of three giant otters averaged as many free-swimming cubs produced per adult group member as larger groups. 10 groups that had free-swimming cubs were recorded again in the following year. Among these, the average cub survival rate after one year was 56%. The mean number of surviving cubs after one year per adult-sized group member was 0.37 (n = 41) (Fig 12).

**Fig 12.**
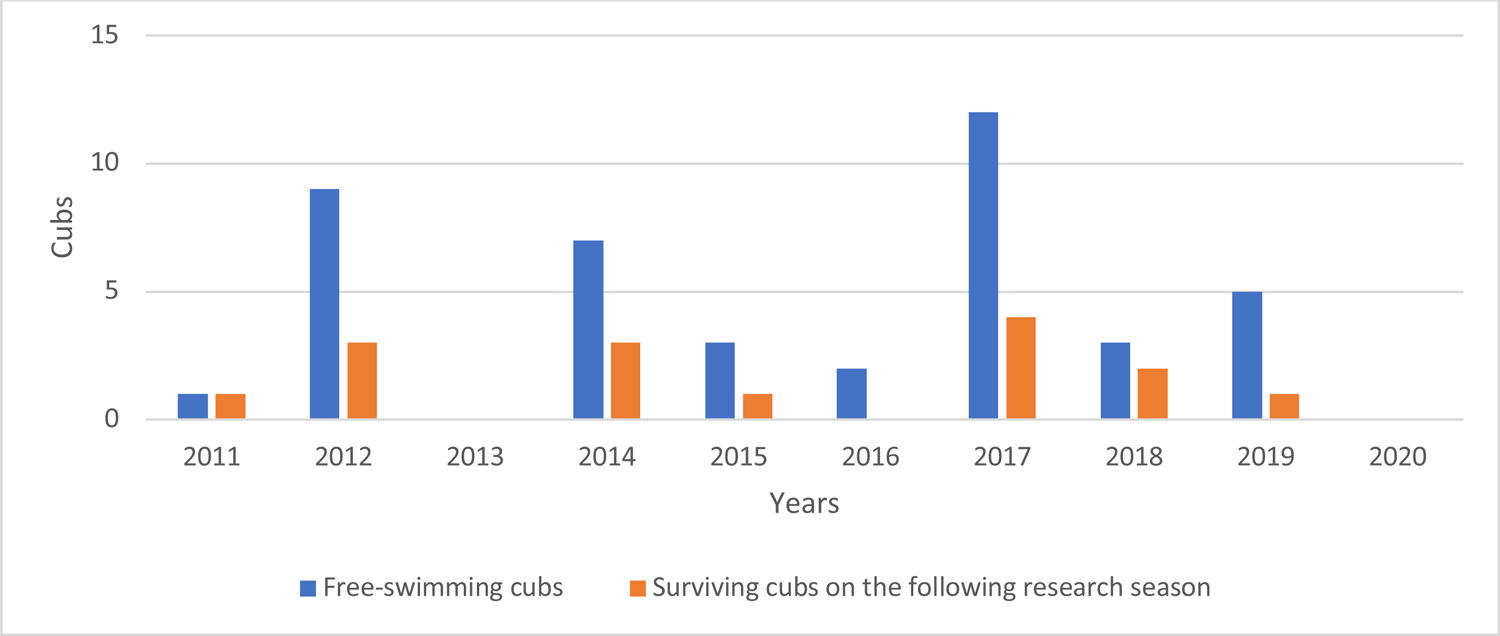
Survival of free-swimming cubs of giant otter after one year in the study area in Cantão, Brazil.

Annual cub production did not seem to correlate with the height or duration of the annual flood but showed a negative correlation with the number of members of immigrant groups that moved into the area during each of these years (r = −0.56) (Fig 13). Data from 2011 and 2012 were excluded from this calculation because part of the study area was not surveyed during those years.

**Fig 13.**
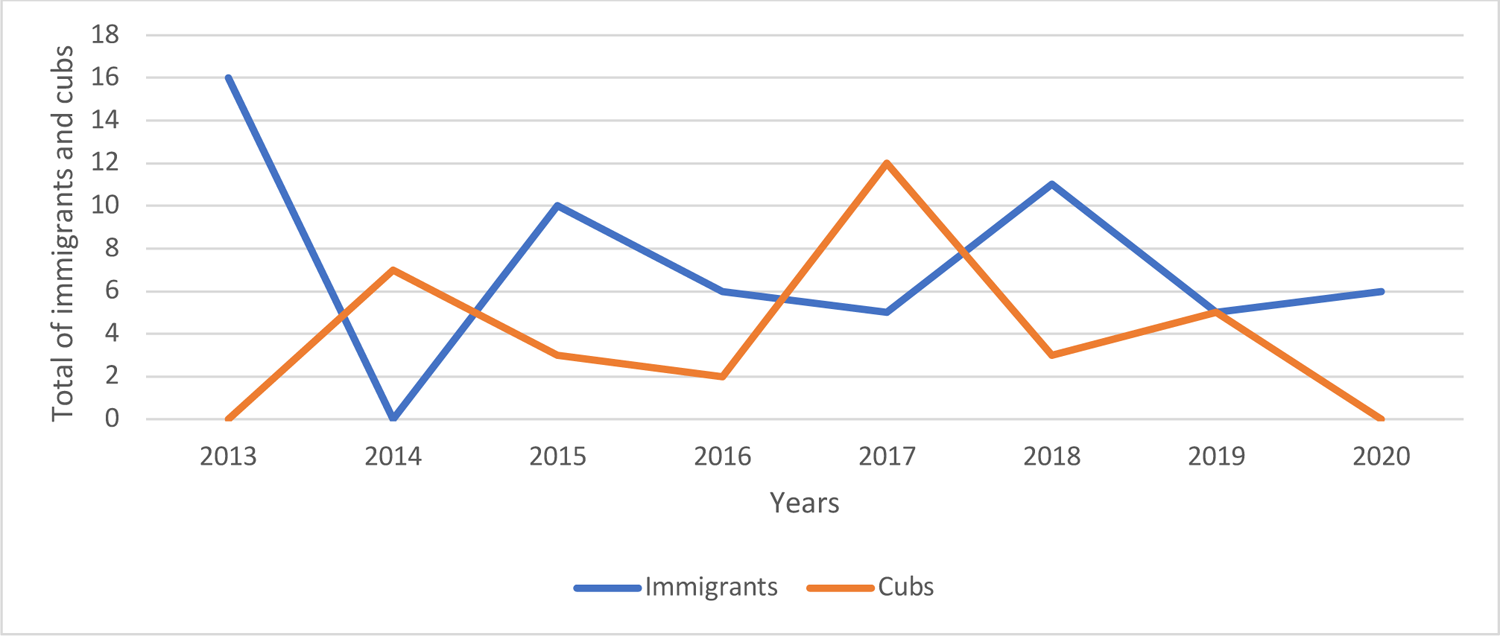
Correlation between number of free-swimming cubs and number of immigrant giant otters in the study area in Cantão, Brazil.

## Discussion

Until recently, giant otters were thought to live in stable family groups occupying stable home ranges [1, 14]. More recent studies showed that groups sometimes include unrelated members [15], and that group home ranges can overlap those of other groups [16, 17]. In Cantão we found giant otter group composition and home ranges to be very fluid. Group home ranges overlap extensively and shift from year to year, with multiple groups sharing the largest lakes, but not synchronically. Even dens are commonly used by different groups. Group composition also changes constantly, not only as cubs are born and adult members leave, but also as new adult-sized individuals join existing groups, eliciting a complex social system in the species.

In the Pantanal, Ribas et al. [15] found that some groups included subordinate individuals that were not offspring of the breeding pair, whereas Leuchtenberger and Mourão [17] also observed adult-sized animals entering new groups as subordinates. In southeast Peru, by contrast, Groenendijk et al [18] found that immigrants were only recruited into a resident group if they claimed the dominant breeding status. In our study we observed adult-sized individuals joining established groups as both subordinate and breeding members. Nine out of 23 resident groups recorded (39%) were observed to accept new members throughout the study. Of 36 adult-sized new group members whose origin was determined during the study, 15 were cubs that had survived their first year while 21 had joined the group as adults. This suggests that giant otter groups in Cantão may, on average, contain more immigrants than offspring of the breeding pair as subordinate members.

Despite sharing most or all of their home range with other groups, giant otter groups in Cantão are quite successful at avoiding one another. Only 16 times during the study we recorded different groups in the same lake on the same day. Agonistic encounters also appear to be rare. While we sometimes saw otters with bite marks from fights that quickly healed, we never saw a seriously injured otter, suggesting that individuals engaged in few agonistic encounters. Agonistic encounters also appear to be uncommon in southeast Peru [18]. These observations contrast with findings in the Pantanal, where agonistic interactions were frequently recorded [17, 19–21].

### Reproduction

Reproduction of giant otters in Cantão showed great annual variability, with resident groups producing free-swimming cubs on average once every three years. This contrasts with what was observed in southeast Peru [18], where resident groups produced one litter per year. Variations in the annual number of cubs produced did not correlate with flood level or duration but showed an inverse relationship with the total number of resident adults and with the number of individuals in groups that immigrated into the area during the previous year. This suggests that successful reproduction of giant otters in Cantão may be depressed by an increase in the density of resident giant otters as well as by the disruptions provoked by the arrival of new individuals or groups into a patch of habitat [18]. Mourão and Carvalho [19], observed infanticide and cannibalism in the species by a solitary male that was located close to the home range of a group formed by six adults and agonistic behaviors were reported in areas with high individual density or in contact zones of two groups territories [21, 22]. Males that enter in a new group, however, often adopt the former youngs and contribute to their raising [9].

In Cantão none of the eleven resident pairs recorded during the study reproduced successfully, and none of the 12 transient pairs observed were accompanied by cubs. In contrast, groups of six or three giant otters were more successful in the number of cubs produced per group-year and in terms of the total number of cubs produced throughout the study. Groups of four and five individuals had a slightly lower reproductive rate than groups of three. This suggests that pairs of giant otters were generally unsuccessful in reproducing in this environment and that the formation of trios of giant otters is critical to the reproduction of the species in Cantão. All six trios whose formation was observed were formed through the recruitment of an adult individual by an existing pair.

The observed cub survival rate after one year (55%, n = 27) is comparable to that reported in other studies [23]. Groenendijk et al. [18] found that 63% of cubs survived their first year in southeast Peru.

### Dispersal and Group Formation

Observed group size in Cantão differs from that reported in other studies. In southeast Peru [18] reported groups of up to 13 individuals, with a mean group size of six otters. In the Xixuaú Preserve in Brazil, the mean group size was 4.5 individuals [3], whereas in the Pantanal mean group size was 4.8 [24]. The maximum group size reported has 15 individuals [25]. However, these studies did not report whether group size counts included cubs or only adults. Our group size counts excluded cubs, as the number of cubs accompanying a group is not stable, varying with new births and cub mortality. In Cantão the maximum group size observed was eight individuals, with a mean and median of four individuals per group. Only four groups with more than six members were recorded during the study and none of them reproduced. Three of these larger groups were reduced to six members in the following year, and one lasted for two years as a resident non-breeding group of eight members before being losing two members, after which the group had a litter. This preponderance of groups of three otters in Cantão further underscores the importance of trios in the ecology of giant otters in the region.

Dispersing giant otters are believed to go through a solitary transient phase during which they search for a mate and an empty patch of territory [2]. In Cantão we recorded 17 transient and 11 resident solitary individuals, with observed periods of residency varying from two months to over a year. Nine of these resident solitaries (82%) eventually formed a pair or joined a resident group; only two transient solitaries (12%) were observed to do so during the same period. Seven out of eight solitaries males recorded were new to the study area, while all solitaries females (N = 4) originated from resident groups.

Pairs of giant otters recorded during the study tended to be transient and unstable and were unable to reproduce successfully. In 95% of recorded pairs with known history (N = 21) at least one member was a former resident of the study area. Ribas et al. [15] reported that newly formed pairs in the Pantanal also tended to be in the vicinity of the territory of at least one of the original groups. Only one of the eleven resident pairs observed in our study remained a pair for longer than one year. Pairs either became a trio through the recruitment of an outside individual, separated, or left the study area. Individual otters often went through two or more pair and/or trio formation attempts before settling into a stable group or disappearing.

In contrast to pairs, trios of giant otters tended to be stable, resident, and to reproduce successfully. Trios accounted for 35% of all resident groups recorded during the study and for 38% of all free-swimming cubs produced. Cub survival after one year for trios was 55% (N = 11), while for pairs it was 0% (N = 2) and for larger groups, it was 64% (N = 14). Once a group became a trio it was able to grow larger by the addition of surviving cubs from previous years as well as immigrant individuals. However, as group size increased, the tendency for members to leave or disappear from the group also increased. Of 14 trios recorded that were seen again in the following year, three (21%) had been reduced to two members, seven (50%) remained as a trio, and four (29%) had increased in size. Of 17 groups of four or more members seen again in the following year, nine (53%) decreased in size, six (35%) remained with the same number of members, and only two (12%) increased in size. By contrast, eight (36%) out of a total of 22 resident and transient pairs recorded became trios during the study. Additional transient pairs may have become trios and left the study area without being recorded as a trio. This suggests that the trio of adult-sized otters is a stable group configuration for giant otters in Cantão.

### Home range shifting and overlap

Home range overlap observed for resident giant otters in Cantão was very common. Most groups shared their home range with at least one other group, and larger lakes were often shared by four or five groups at different times. Groups rarely used the same lake for more than a few days, and when they left, they were often replaced by other groups. Some groups used certain lakes for alternating periods of one to several days over a month or more, often sleeping in the same dens and using the same latrines used previously by other groups. Groups with small cubs sometimes remained in the same lake for a month or more, but generally moved the cubs to a different lake at least once before they became free-swimming. This contrasts with findings by Staib [13] that indicate that giant otter ranges do not overlap at all in oxbow lake environments in southeast Peru. Evangelista and Rosas [26] observed partial range overlap in a tropical river habitat. Leuchtenberger et al. [25] also observed partial range overlap along linear river habitat in the Pantanal. The home range of some groups in the Pantanal overlapped partially with that of neighboring groups, but each group appeared to have a core territory that is actively defended. Although groups in the Pantanal tended to not use overlapped areas at the same time, 12 agonistic encounters between groups were observed over a two-year study, including fights [27]. In Cantão only three agonistic encounters were recorded over ten years, both limited to territorial vocalizations between groups, but without fights.

Almost all observed groups shifted at least part of their home range from year to year, and many shifted to a completely different set of lakes between years. The high turnover rate of resident groups within the core study area is indicative of the frequency of large-scale shifting of home ranges. Sometimes group home ranges drifted slowly over the years until they left the study area, and other times a group would move to a completely new home range from one year to the next. Only seven of 23 (30%) resident groups remained in the study area for more than two years, and only five (21.7%) remained for more than three years. Regardless of how much they shifted their home ranges, groups were faced with a new set of neighbors each year, often sharing some of the same lakes. In oxbow lakes in southeast Peru, resident groups tend to remain within the same home range indefinitely [18, 28]. In the continuous river habitat of the Pantanal, Leuchtenberger et al. [27] observed shifting home ranges in a pattern similar to that observed in Cantão.

### Plasticity of Giant Otter Social and Territorial Behavior

The observed differences in giant otter group dynamics and territorial behavior between Cantão and other sites can be explained by the spatial characteristics of the habitat. Previous long-term studies of the species were conducted in areas composed of patchy (isolated oxbow lakes) or linear (rivers) habitats. In the Cantão flooded forest ecosystem, as in some parts of the Brazilian Pantanal [17], optimal giant otter habitat is continuous in all directions. In the dry season, most of Cantão lakes are connected within a few hundred meters of several other lakes. We also observed that giant otters use the flooded forests during the flood season. This affects both dispersal opportunities and cost-benefit tradeoffs for territorial defense.

Every resident giant otter group in our study had several other groups residing within a few hundred meters of its home range, and most of them shared part or all of their home range with up to six other groups. High-quality habitats can favor individual propensity to emigrate [29]. Dispersing giant otters in Cantão not only have a hospitable and predictable environment in all directions, which they may explore before emigrating, they are also familiar with potential partners in the surrounding area, some of which may be scent-marking at the same sites as the potential disperser’s group. In the isolated oxbow lake environments studied elsewhere, potential dispersers may have to transit large patches of a suboptimal environment with which they are unfamiliar and depend on chance to meet potential partners. In linear river habitats, the potential disperser may find optimal habitat extending in one dimension, and maybe familiar with potential partners belonging to upstream and downstream neighboring groups. This habitat effect can explain the observed increase in average group size as individuals move from continuous and bidimensional habitats (flooded forest with high oxbow lake density) to linear but continuous habitats (rivers), to patchy discontinuous habitat (isolated lakes).

The fact that optimal habitat in Cantão is continuous but also fragmented into individual lakes may explain the tolerance for home range overlap displayed by resident giant otter groups. A “dear enemy” effect [30], where territorial animals direct less aggression toward established neighbors than toward strangers, maybe at play. Although the dear enemy effect is more common when neighbors had well-established territories [31–33], the use of scent cues for individual and group recognition may act as a way to reduce aggressiveness in these fluctuating territories [34–36]. The resource availability in Cantão also renders the circumstantial benefits toward aggressive behaviors between groups to be minimal. Fish prey is abundant in hundreds of Cantão lakes, but foraging giant otters are constantly on the move, rarely stopping for more than a few minutes even at the most productive sites, probably reducing disturbance effects of fishing on fish wariness. Foraging giant otter groups create considerable disturbance through splashing, jumping, and turbulent swimming, and groups are soon forced to move to a different lake to continue foraging, even if the lake they just traversed still has plenty of fish. If a group arrives at a lake and finds that another group is already there, it may derive little immediate benefit from chasing the other group away because the lake has been disturbed, providing few foraging returns, being more profitable just to move on to another lake. A group wishing to avoid conflict can easily avoid encountering other groups by simply moving to one of many nearby lakes. Since lakes in Cantão are not large enough to be occupied continuously by a single group, a group cannot secure exclusive use of a lake no matter how much energy is expended in territorial defense. The optimal solution appears to be to tolerate other resident groups sharing the lakes within its home range as long as fish prey availability does not become a limiting factor.

The “dear enemy” effect is facilitated by the ability to recognize familiar neighbors [37], and giant otters are particularly well adapted for this due to their individual throat markings, scent cues, familiar sounds, and periscoping behavior. This may also explain the tendency for the total number of adult otters in our research area to remain within a relatively narrow range even with a high annual turnover of resident groups.

The high annual variation in the number of cubs produced by giant otter groups in Cantão is also different from what was reported from other sites. This may also be explained by the specific territorial dynamics generated by the local landscape. We found a negative correlation between production of free-swimming cubs and the number of new adults moving into the area. A high proportion of newly arrived groups increases the likelihood of stressful encounters and/or costly avoidance behavior between groups. In captivity, stress caused by visitors can cause a giant otter mother to stop lactating [38]. Londoño and Tigreros [39] reported that stress caused by noise or the presence of strangers caused giant otters to carry pups under 30 days old into the water, where in the wild they would be at risk of drowning or encountering predators. Schenk and Staib [40] observed that reproductive success was depressed for giant otters living in lakes heavily visited by tourists. Likely, increased stress caused by increased population density or the arrival of unfamiliar groups depresses the reproductive rate of giant otters in Cantão, and this may also contribute towards the maintenance of population density close to the environment’s carrying capacity.

### Implications for Conservation

The main bottleneck for successful colonization of new habitat by dispersing giant otters is whether a dispersing individual can meet a potential mate at the right time, in a suitable place [28]. If so, colonization of new areas may be more difficult in environments like Cantão, where it appears that the formation of a trio of giant otters is a prerequisite for successful reproduction. This possibility should be taken into account in reintroduction projects for the species, which currently assume that the introduction of pairs of animals into the unoccupied habitat is sufficient to start a new population [41]. Our findings also indicate that giant otters may change partners several times before settling into a stable group and successfully reproducing. This may be due to genetic or other incompatibilities that require trial and error to avoid. Isolated reintroduced groups may not be able to reproduce successfully even if they were captured and relocated as a group.

If the hypothesis of depressed reproductive success caused by stress provoked by encounters with strangers is correct, it could mean that frequent encounters with humans may also reduce the rate of reproduction of giant otters. Even when intruding humans don’t directly encounter giant otter groups, the disturbance of prey by fishing or other activity may have an effect analogous to an additional giant otter group foraging through the habitat, and if it occurs repeatedly, it may reduce reproductive success and decrease the area’s carrying capacity. We documented three episodes of giant otters relocating very young pups after brief encounters with intruders in Cantão. Two of these episodes were merely a motorboat passing by the breeding den. The same groups were largely indifferent to approaches by researchers with whom they were familiar, to the extent of bringing out the cubs for swimming lessons in the presence of five researchers observing without cover from less than 100 meters away. Breeding refuges where humans are excluded, or allowed only under strict regulation and monitoring, may be essential to the reproduction of giant otters. This reinforces the importance of the strict protected areas (IUCN Category 2 or higher) with zones where no visitation is allowed for the conservation of the species.

The Cantão ecosystem appears to sustain a high density of giant otters compared to other sites, mainly due to its abundance of fish prey and suitable habitat. Overall giant otter densities at other protected areas tend to be relatively low because these areas consist largely of unsuitable habitat, while all of Cantão State Park consists of habitat similar to that of the study area. If the density of resident groups in the 16 lakes of our study area is indicative of the density of the species throughout the park’s 850 oxbow lakes, this may be one of the most important protected areas for the species today. Habitat similar to Cantão’s, with large numbers of oxbow lakes within an igapó flooded forest matrix, also occurs at other sites in the Amazon basin, such as along the lower reaches of the Juruá, Purus, Tefé, and Jaú rivers. Identifying and surveying these sites, even if giant otters are currently rare or absent in some of them, may help to identify critical areas for the recovery and protection of the species.

## Acknowledgments

The authors would like to thank Dr Nicole Duplaix and Dr Christof Schenk for their support and technical advice throughout the years. Special thanks to Dr Robin Williams and Ms. Daniella Woodall for providing technical assistance in the field, and to Ms. Cheryl Williams for her support. We also extend our gratitude to our volunteers Adriana Luz, Melissa Savage, Jean-Paul Magnan, Elena Weindel, Sarah Griswold, Iris Berger and Gabriel Mihahira for their contribution to this research. We are in debt to all the staff and rangers of Instituto Araguaia over the years, in particular to our deceased ranger Mr. Manoel Dias da Silva, native of Cantão and dedicated conservationist, for his intrinsic knowledge and precise insights on giant otters and their habitat.

